# Mitochondrial Network Configuration Influences Sarcomere and Myosin Filament Structure in Striated Muscles

**DOI:** 10.1101/2022.01.14.476070

**Authors:** Prasanna Katti, Alexander S. Hall, Peter T. Ajayi, Yuho Kim, T. Bradley Willingham, Christopher K. E. Bleck, Han Wen, Brian Glancy

## Abstract

Sustained muscle contraction occurs through interactions between actin and myosin filaments within sarcomeres and requires a constant supply of adenosine triphosphate (ATP) from nearby mitochondria. However, it remains unclear how different physical configurations between sarcomeres and mitochondria alter the energetic support for contractile function. Here, we show that sarcomere cross-sectional area (CSA) varies along its length in a cell type-dependent manner where the reduction in Z-disk CSA relative to the sarcomere center is closely coordinated with mitochondrial network configuration in flies, mice, and humans. Further, we find myosin filaments near the sarcomere periphery are curved relative to interior filaments with greater curvature for filaments near mitochondria compared to sarcoplasmic reticulum. Finally, we demonstrate smaller myosin filament lattice spacing at filament ends than filament centers in a cell type-dependent manner. These data suggest both sarcomere structure and myofilament interactions are influenced by the location and orientation of mitochondria within muscle cells.

## Introduction

Force generation within striated muscle cells occurs through the formation and release of bonds between actin and myosin filaments within the sarcomere with networks of highly connected sarcomeres^1–4^ occupying up to 85% of cellular volume^4–6^. Sustained muscle contractions also necessitate continued provision of calcium from the sarcoplasmic reticulum (SR) and adenosine triphosphate (ATP) from mitochondria and/or glycolytic enzymes, thereby requiring significant cellular volume in proximity to the sarcomeres to be invested in these systems as well^7^. The chronic force production demand (i.e., magnitude and frequency) on a given muscle cell type is well known to influence the amount of cellular volume allocated to, and thus the functional capacity of, each component^7–16^. However, it is less well understood how variations in contractile demand affect the spatial organization among the tightly packed sarcomeres and the organelles that must fit in between them or how different cellular architectures alter the functional support for muscle contraction. In particular, while mitochondria are known to form cell-type dependent networks aligned perpendicular and/or in parallel to the contractile axis of the muscle cell^11,17–20^, the functional implications of placing a mitochondrion in parallel between sarcomeres versus wrapping a mitochondrion perpendicularly between sarcomeres near the Z-disks remains unclear.

Here, we investigate the detailed physical interactions among sarcomeres and their adjacent organelles in eleven muscle types across three species (human, mouse, fly) using high resolution, 3D volume electron microscopy. We demonstrate that sarcomere cross-sectional area (CSA) is smaller at the Z-disk ends than in the sarcomere centers where myosin resides. Further, we find that the magnitude of intrasarcomere CSA heterogeneity is cell type-dependent and closely coordinated with the location of mitochondria between the sarcomeres rather than the total volume or size of mitochondria within the muscle cell. By performing a massively parallel myosin filament analysis, we show that intrasarcomere CSA heterogeneity is achieved, at least in part, by curvature of the contractile filaments near the periphery, but not the center, of each sarcomere. Across muscle cell types, the magnitude of myosin curvature is highest in cells with large proportions of mitochondria running perpendicularly near the Z-disk, while within cells, myosin curvature is greater for filaments near mitochondria and lipid droplets compared to the sarcoplasmic reticulum. Moreover, intrasarcomere heterogeneity in myosin shape results in variable filament-to-filament lattice spacing along the length of the sarcomere where myosin filaments are closer together near the filament ends than in the middle. Together, these data indicate that sarcomere and myosin filament structure are influenced by where mitochondria are placed within a muscle cell.

## Results

### Sarcomere Cross-Sectional Area Varies Along its Length in a Cell Type-Dependent Manner

To better understand how muscle cell architecture relates to muscle cell function, we performed a 3D survey of sarcomere and adjacent organelle structures across eleven muscle types whose contractile functions have generally been well defined^1,21–25^. We performed focused ion beam scanning electron microscopy (FIB-SEM) with 5-15 nm pixel sizes on cardiac and skeletal muscle tissues fixed *in vivo* in mice and on skeletal muscle tissues from *Drosophila melanogaster* (18 FIB-SEM datasets available at doi.org/10.5281/zenodo.5796264). Close inspection of a mouse fast-twitch glycolytic muscle raw FIB-SEM volume (**Figure 1a**, **Supplementary Video 1**) suggested that sarcomere cross-sectional area (CSA) was variable at different regions along its length with the CSA becoming smaller near the Z-disk ends compared to the center of the sarcomere within the A-band (myosin containing region). Indeed, 3D rendering of the fast-twitch glycolytic muscle sarcomeres revealed large gaps between the parallel sarcomeres near the Z-disks (**Figure 1b**, **Supplementary Video 2**) and these gaps corresponded to where perpendicularly aligned mitochondria (**Figure 1c-e,g-h**) and SR doublets (**Figure 1d,h**) were located adjacent to the sarcomere. Conversely, *Drosophila* indirect flight muscles (IFM) appear to maintain a constant CSA across the entire sarcomere with no large gaps apparent between parallel sarcomeres (**Figure 1i**, **Supplementary Video 3**) despite a large volume of adjacent mitochondria (**Figure 1j**). Thus, we hypothesized that intrasarcomere CSA heterogeneity was cell type-dependent. To test this hypothesis, we compared the maximal CSA at the Z-disk and corresponding A-band for each half sarcomere sheet (**Supplementary Video 4**) within muscle volumes from six mouse muscle types and four *Drosophila* muscle types, as well as in human muscles previously published as part of the Baltimore Longitudinal Study of Aging^19^. The magnitude of intrasarcomere CSA heterogeneity was highly variable across cell types (**Figure 1k**) with the lowest values observed in *Drosophila* IFM and jump muscles (4.7±2.4% CSA difference between A-band and Z-disk, n=3 cells, 56 half sarcomere sheets and 9.5±0.8% difference, n=3 cells, 84 half sarcomere sheets for IFM and jump muscles, respectively) and mouse cardiac muscles (10.9±1.8% difference, n=5 cells, 122 half sarcomere sheets). The greatest degree of intrasarcomere CSA heterogeneity was found in the *Drosophila* leg muscles (40.3±6.5% difference, n=5 cells, 21 half sarcomere sheets), mouse oxidative muscles (34.4±4.8% difference, n=4 cells, 86 half sarcomere sheets, and 31.1±1.7% difference, n=4 cells, 84 half sarcomere sheets for fast- and slow-twitch muscles, respectively) and human muscles (29.3±1.6% difference, n=3 cells, 80 half sarcomere sheets). The wide range of intrasarcomere CSA heterogeneity values across cell types in both fly and mouse muscles, as well as the high values observed in humans, suggest that cell type specific intrasarcomere CSA heterogeneity is conserved from invertebrate to human muscles.

**Figure 1:**
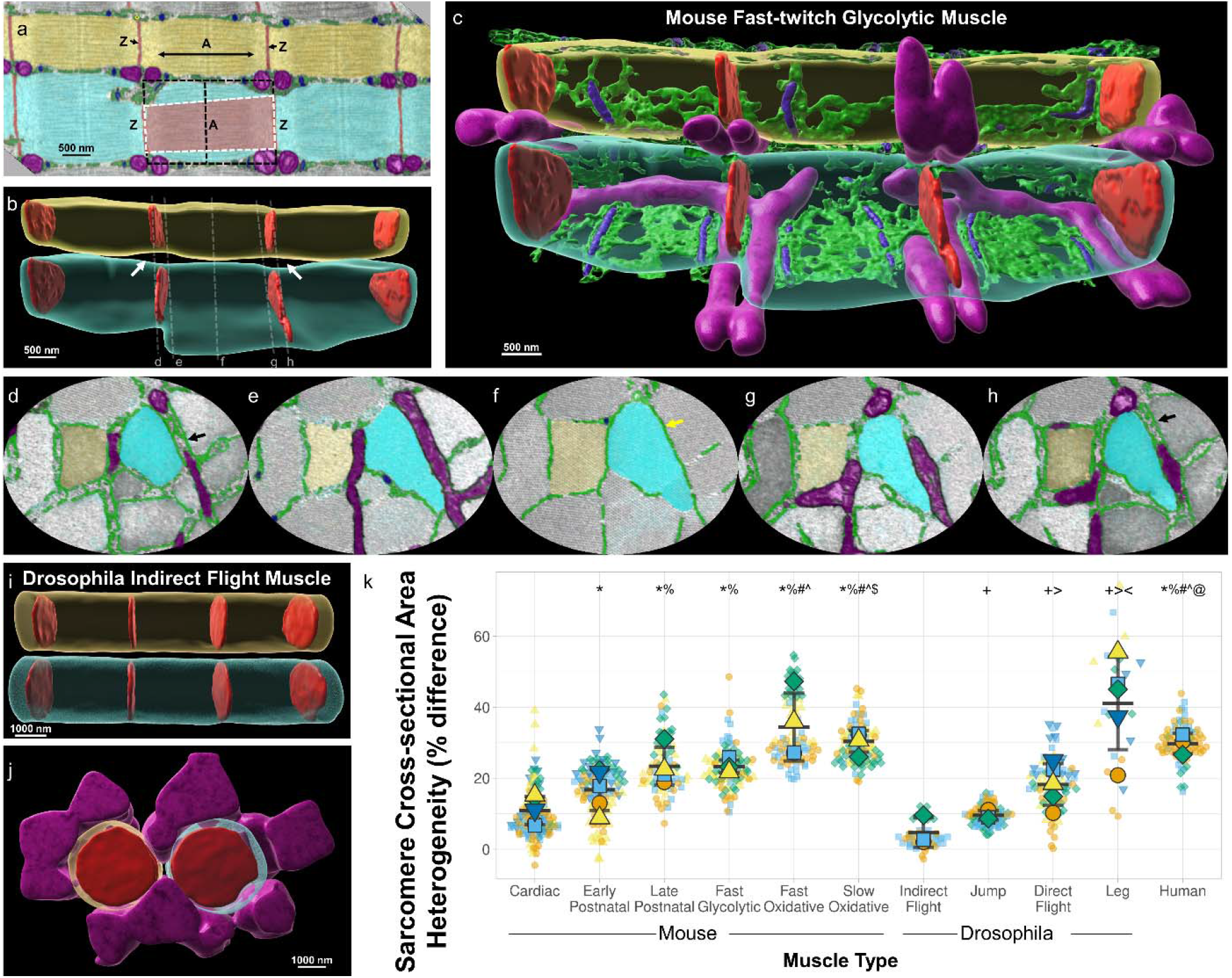
Intrasarcomere Cross-sectional Area Heterogeneity is Dependent on Muscle Cell Type. a) Longitudinal raw focused ion beam scanning electron microscopy (FIB-SEM) image of a mouse fast-twitch glycolytic gastrocnemius muscle showing two parallel myofibrillar segments comprised of three sarcomeres each (cyan and yellow). Diameter of the A-band (A) is larger than the diameter of the Z-disk (Z, red). Mitochondria (magenta), sarcoplasmic reticulum (SR, green), and t-tubules (T, blue) are also shown. b) 3D rendering of the sarcomere boundaries from a (translucent cyan and yellow) and their Z-disk structures (red). White arrows highlight gaps between in parallel sarcomeres. c) Same 3D rendering as b but also showing sarcomere adjacent mitochondria (magenta), and partial sarcoplasmic reticulum (green) and t-tubule (blue) structures. d-h) Raw FIB-SEM cross-sectional views at the Z-disk, I-band, and A-band regions indicated by dotted lines in b highlighting the varying diameters of the mitochondrial and SR/T structures located between in parallel sarcomeres (cyan and yellow). Black arrows highlight SR doublets. Yellow arrow highlights longitudinal SR. i) 3D rendering of the sarcomere boundaries (translucent cyan and yellow) for six sarcomeres from *Drosophila* indirect flight muscle and their Z-disk structures (red). j) 90° rotation of the indirect flight muscle sarcomeres from i also showing adjacent mitochondria which do not run perpendicular to the contractile axis. k) Percent difference in sarcomere cross-sectional area at the A-band relative to the Z-disk per half sarcomere sheet across eleven muscle cell types. Large shapes represent individual cell values, smaller shapes represent values per individual half sarcomere sheet. Black lines represent cell Mean±SD. *significantly different from Cardiac. %significantly different from Early Postnatal (postnatal day 1). #significantly different from Late Postnatal (postnatal day 14). ^significantly different from Fast Glycolytic. $significantly different from Fast Oxidative. @significantly different from Slow Oxidative. ^+^significantly different from Indirect Flight. >significantly different from Jump. <significantly different from Direct Flight. N values: Cardiac-5 cells, 122 half sarcomere sheets; Early Postnatal-5 cells, 106 half sarcomere sheets; Late Postnatal-4 cells, 70 half sarcomere sheets; Fast Glycolytic-4 cells, 86 half sarcomere sheets; Fast Oxidative-4 cells, 86 half sarcomere sheets; Slow Oxidative-4 cells, 84 half sarcomere sheets; Indirect Flight-3 cells, 56 half sarcomere sheets; Jump-3 cells, 84 half sarcomere sheets; Direct Flight-5 cells, 80 half sarcomere sheets; Leg-5 cells, 21 half sarcomere sheets; Human-3 cells, 80 half sarcomere sheets.

### Sarcomere Shape is Coordinated with Mitochondrial Location

The six muscle types with greater than 20% intrasarcomere CSA heterogeneity in Figure 1 are all known to have large proportions of their mitochondrial networks which wrap perpendicularly around the sarcomere adjacent to the Z-disks, whereas the five muscle types with less than 20% heterogeneity all have mitochondrial networks primarily oriented in parallel to the contractile apparatus^11,18,19,26–29^. Thus, we hypothesized that sarcomere structure is linked to mitochondrial network configuration. To test this hypothesis, we first used machine learning^30^ to segment out mitochondrial network structures within each FIB-SEM dataset^11^ (**Supplementary Video 5**). To assess mitochondrial network configuration, we further separated the mitochondrial network structures into regions adjacent to the Z-disk (light blue in **Supplementary Videos 6-7**) and regions not adjacent to the Z-disk (red in **Supplementary Videos 6-7**). Using a distance from Z-disk threshold of 200 nm longer than the width of the sarcomeric I-band (actin only region) in each dataset resulted in all perpendicularly oriented mitochondria to be included in the Z-adjacent mitochondrial pool (**Figure 2a-k, Supplementary Video 6**). We then compared the percentage of the total mitochondrial network that is adjacent to the Z-disk versus the magnitude of intrasarcomere CSA heterogeneity for each dataset which revealed a significant linear relationship (**Figure 2l**, R^2^=0.4964, p<0.001) indicating that greater intrasarcomere CSA heterogeneity is associated with more of the mitochondrial pool being localized to the Z-disk. Comparing intrasarcomere CSA heterogeneity to total mitochondrial content also revealed a statistically significant linear relationship (**Figure 2m**, R^2^=0.1076, p=0.024), albeit negative and much weaker than with Z-adjacent mitochondrial abundance. Indeed, in a two component multiple regression model evaluating the impact of both Z-adjacent mitochondrial abundance and mitochondrial content on intrasarcomere CSA heterogeneity, Z-adjacent mitochondrial abundance contributes significantly (standardized beta coefficient 0.709, p<0.001) whereas mitochondrial content does not (standardized beta coefficient −0.005, p=0.968). Additionally, there was no significant correlation between intrasarcomere CSA heterogeneity and individual mitochondrial volume (**Figure 2n**, R=0.059, p=0.126). These data suggest that sarcomere shape appears to be closely coordinated with where mitochondria are located within a muscle cell, and less so with how many or how big the mitochondria are.

**Figure 2:**
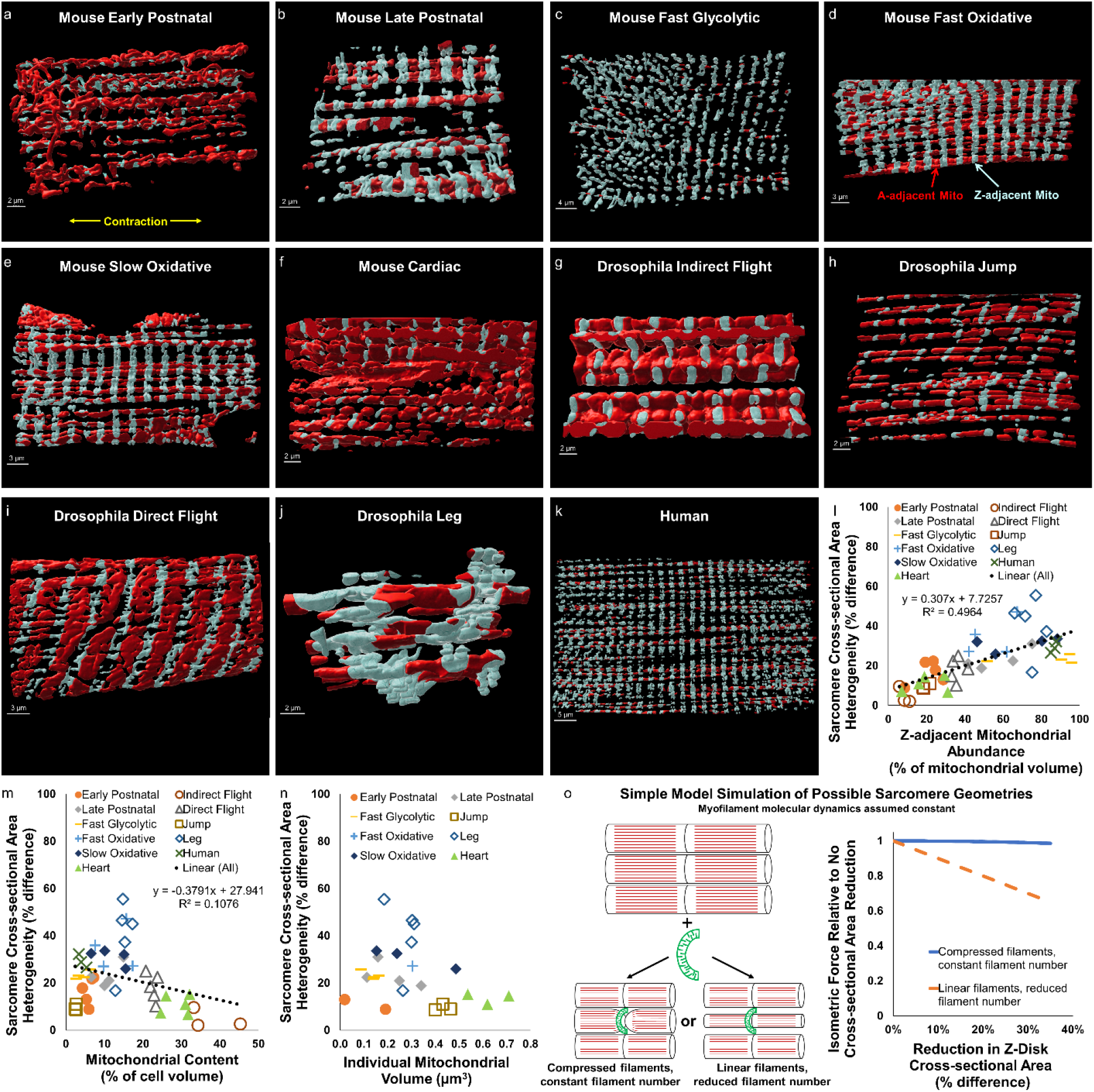
Magnitude of Intrasarcomere Cross-sectional Area Heterogeneity is Related to Mitochondrial Location. a-k) 3D rendering of representative mitochondrial network structures from eleven muscle types from mouse, *Drosophila,* and human tissues showing Z-adjacent regions (light blue) and A-adjacent regions (red). l) Linear correlation (black dotted line, p<0.001) between Z-adjacent mitochondrial abundance and sarcomere cross-sectional area (CSA) heterogeneity. Individual points represent individual muscle cell values. m) Linear correlation (black dotted line, p=0.024) between mitochondrial content and sarcomere CSA heterogeneity. n) Lack of correlation between individual mitochondrial volume and sarcomere CSA heterogeneity. o) Simulations of simple sarcomere geometry models showing proportional reduction in isometric force when entire sarcomere CSA is reduced and negligible reduction in force when only the Z-disk CSA is reduced.

However, closer inspection of **Figure 2m** suggests that the intrasarcomere CSA heterogeneity/mitochondrial content relationship may actually be biphasic in nature with a transition occurring between 14.5-17.5% mitochondrial content. Indeed, separating the data into high (>14.5%) and low (<17.5%) mitochondrial content groups where both contain the apparent transition region results in a strong positive correlation (R^2^ = 0.409, p<0.001) with intrasarcomere CSA heterogeneity in low mitochondrial content cells (**Supplementary Figure 1a**) and a strong negative correlation (R^2^ = 0.672, p<0.001) with intrasarcomere CSA heterogeneity in high mitochondrial content cells (**Supplementary Figure 1b**). Moreover, both mitochondrial content (standardized beta coefficient 0.521, p<0.001) and mitochondrial location (standardized beta coefficient 0.484, p<0.001) contribute significantly to the prediction of intrasarcomere CSA heterogeneity in a multiple regression model of low mitochondrial content muscle cells, though the contribution of mitochondrial content (standardized beta coefficient −0.238, p=0.185) does not reach significance compared to mitochondrial location (standardized beta coefficient 0.699, p<0.001) in high mitochondrial content cells. These data suggest there is a transition in striated muscle cellular design principles which occurs when mitochondrial content within the intrafibrillar space reaches 14.5-17.5% where additional mitochondria no longer result in intrasarcomere CSA heterogeneity. Indeed, below this threshold, mitochondrial content and mitochondrial location remain independent of one another (**Supplementary Figure 1c**, R^2^=0.034, p=0.314) as mitochondrial networks of similar total content can be primarily parallel (**Figure 2a,h**), grid-like (**Figure 2b,d,e,j**), or perpendicular (**Figure 2c,k**) in nature, as also recently reported in shrew muscle^31^. However, above this threshold, mitochondrial content and location are closely related (**Supplementary Figure 1d**, R^2^=0.687, p<0.001) as additional mitochondria are placed in parallel to the contractile axis and therefore result in a uniform reduction of the entire sarcomere CSA. Thus, while mitochondrial location maintains a consistent relationship with intrasarcomere CSA heterogeneity across all striated muscle cells (**Figure 2l**), mitochondrial content appears to play a secondary role by mediating mitochondrial location once mitochondrial content reaches the 14.5-17.5% cellular design transition point.

**Supplementary Figure 1:**
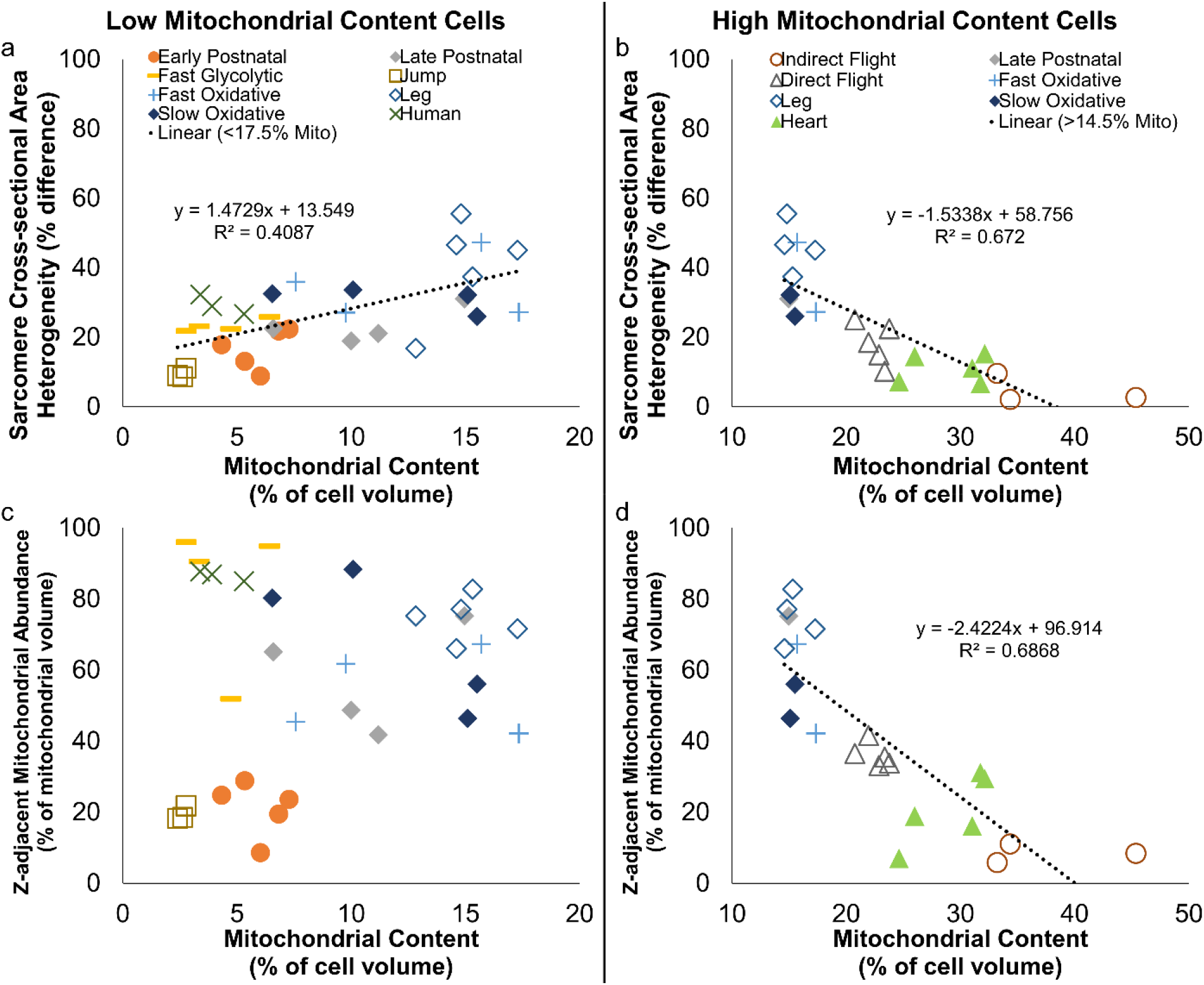
Relationships between Mitochondrial Content, Location, and Sarcomere Shape. a) Correlation (black dotted line, p<0.001) between mitochondrial content and sarcomere cross-sectional area (CSA) heterogeneity for low mitochondrial content (<17.5% volume) cells. Individual points represent individual muscle cell values. b) Correlation (black dotted line, p<0.001) between mitochondrial content and sarcomere cross-sectional area (CSA) heterogeneity for high mitochondrial content (>14.5% volume) cells. c) Lack of correlation between mitochondrial content and Z-adjacent mitochondrial abundance for low mitochondrial content (<17.5% volume) cells. d) Correlation (black dotted line, p<0.001) between mitochondrial content and Z-adjacent mitochondrial abundance for high mitochondrial content (>14.5% volume) cells.

Since there are currently no methods to directly assess forces at the sarcomere scale within intact muscle cells, as a first step toward assessing the functional impact of intrasarcomere CSA heterogeneity, we developed a simple geometrical model to simulate the relative change in force production due to the addition of a new mitochondrion wrapping perpendicularly around the sarcomere near the Z-disk (**Figure 2o**). Simulations of two types of sarcomeric adaptations to mitochondrial addition were performed. First, sarcomere CSA was maintained constant along its length, as is commonly assumed^32–36^, and, thus, reduction of the Z-disk CSA to make space for the mitochondrion results in a proportional loss in the number of linear myofilaments throughout the sarcomere. As expected, simulations of this uniform sarcomere CSA model showed that isometric force production decreased linearly with the magnitude of reduction of the Z-disk CSA (orange dotted line in **Figure 2o**). The second model tested was based on the intrasarcomere CSA heterogeneity structures described above where the Z-disk, but not the middle, of the sarcomere is compressed to make room for the new mitochondrion. It was assumed in the heterogeneous CSA model that myofilaments curved slightly in proportion to the reduction in Z-disk CSA but that filament number remained constant. These simulations revealed negligible loss of force in response to up to 35% reduction of the Z-disk CSA (blue line in **Figure 2o**). Thus, the data from these simulations suggest that by limiting the loss of sarcomere CSA to near the Z-disk, there is little contractile cost to adding a mitochondrion wrapped perpendicularly around a sarcomere. However, it is important to note that this simple geometrical model does not account for any potential changes in myofilament molecular dynamics that may occur within the sarcomere as a result of intrasarcomere CSA heterogeneity. More detailed models accounting for the true 3D myofilament structures and their dynamic molecular interactions within the sarcomere will be needed to more fully understand the functional impact of intrasarcomere CSA heterogeneity.

### Proximity to Mitochondria Influences Curvature of Myosin Filaments

To better understand the influence of intrasarcomere CSA heterogeneity on the myofilaments within the sarcomere, we performed a massively parallel segmentation of all the myosin filaments within a FIB-SEM dataset (**Figure 3a**, **Supplementary Video 8**) which allowed for assessment of hundreds of thousands to millions of individual filaments per cell. Based on electron microscopy images such as **Figure 1a** here as well as in the literature (e.g. Figure 1a in Wang et al.^37^), we hypothesized that intrasarcomere CSA heterogeneity also leads to variability in filament structures within a single sarcomere where filaments near organelles at the sarcomere periphery are slightly curved while those in the center remain linear. To test this hypothesis, we fit each segmented myosin filament to a straight line and then measured the deviation of each filament from linearity as a proxy for filament curvature (**Supplementary Video 9**). Indeed, a large variation in myosin filament curvature can be observed within a single sarcomere (**Figure 3b,c**). To quantify myosin filament curvature relative to intrasarcomere positioning, we performed machine learning segmentation of the major organelles which surround and provide the boundaries for the sarcomeres (sarcoplasmic reticulum + t-tubules, SR/T; mitochondria, Mito; and lipid droplets, LD) (**Figure 3d, Supplementary Video 10**) and then measured the shortest distance between every myosin filament and each organelle. In both slow oxidative and fast glycolytic muscles from the mature mouse, myosin filaments within 100 nm of the overall sarcomere boundary are significantly less linear than those more toward the center (**Figure 3e**). Proximity to each individual organelle (SR/T, Mito, LD) also corresponds to greater myosin filament curvature than more distal filaments in slow oxidative and fast glycolytic muscles (**Supplementary Figure 2a-d**). However, proximity to mitochondria and lipid droplets results in greater myosin filament curvature than proximity to SR/T or the overall sarcomere boundary (**Figure 3f**). These data demonstrate that myosin filament structure is heterogeneous within sarcomeres with slightly curved filaments near the periphery and more linear filaments within the core of the sarcomere. Additionally, proximity to larger organelles such as mitochondria and lipid droplets is associated with greater myosin filament curvature than smaller diameter organelles such as the SR/T.

**Figure 3:**
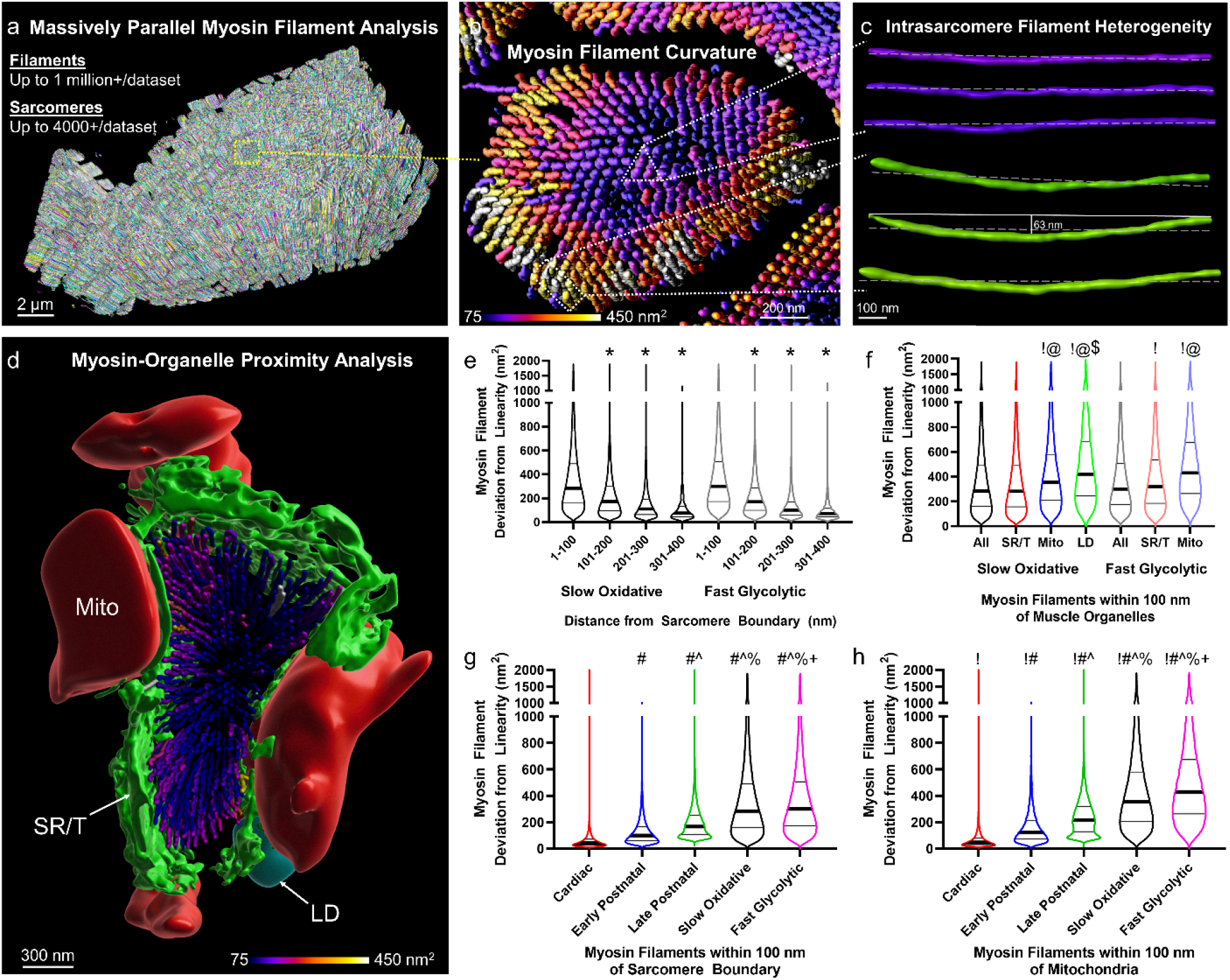
Intrasarcomere Myosin Filament Heterogeneity and Organelle Proximity. a) 3D rendering of 364,000+ myosin filaments (various colors) from a mouse late postnatal muscle. b) 3D rendering of the myosin filaments from the highlighted sarcomere in a. Filament color represents filament deviation from linearity. c) 3D rendering of three myosin filaments from the sarcomere core in b (purple, upper) and three filaments from the periphery (green, lower) showing the variability in filament linearity. d) 3D rendering of the myosin filaments from a single mouse cardiac sarcomere and the organelles which comprise the sarcomere boundary. Mitochondria (Mito, red), sarcotubular network (SR/T, green), and lipid droplets (LD, cyan) are shown. e) Myosin filament deviation from linearity as a function of distance from the sarcomere boundary for Slow Oxidative and Fast Glycolytic fibers. f) Myosin filament deviation from linearity for filaments within 100 nm of different muscle organelles. g) Myosin filament deviation from linearity for filaments within 100 nm of the sarcomere boundary across cell types. h) Myosin filament deviation from linearity for filaments within 100 nm of mitochondria across cell types. Thick black lines represent median values, thin black lines represent upper and lower quartile values. Width of the violin plot represents the relative number of filaments at a given value. *significantly different from 1-100 nm. ǃsignificantly different from All. @significantly different from SR/T. $significantly different from mitochondria. #significantly different from Cardiac. ^significantly different from Early Postnatal. %significantly different from Late Postnatal. “significantly different from Slow Oxidative. N values: Slow Oxidative-700,950 filaments; Fast Glycolytic-680,038; Cardiac-265,340; Early Postnatal-463,176; Late Postnatal-364,494.

**Supplementary Figure 2:**
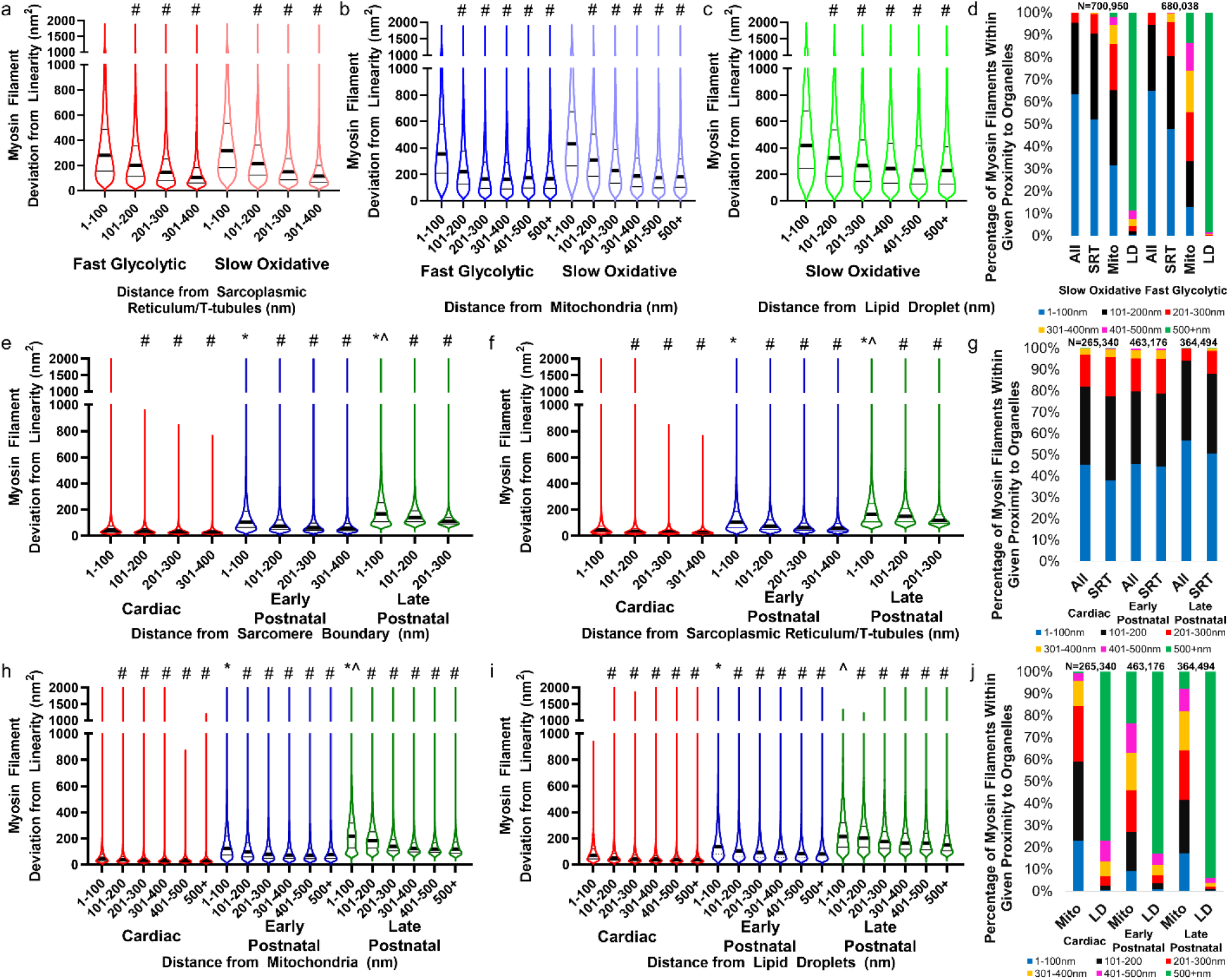
Myosin filament curvature and organelle proximity in mouse muscles. a-c) Myosin filament deviation from linearity as a function of distance from sarcoplasmic reticulum/t-tubules (a), mitochondria (b), and lipid droplets (c) for Slow Oxidative and Fast Glycolytic fibers. d) Percentage of myosin filaments within 100 nm distances from sarcomere boundary organelles in Slow Oxidative and Fast Glycolytic muscles. e,f,h,i) Myosin filament deviation from linearity as a function of distance from the sarcomere boundary (e), sarcoplasmic reticulum/t-tubules (f), mitochondria (h), and lipid droplets (i) for Cardiac, Early Postnatal, and Late Postnatal fibers. g,j) Percentage of myosin filaments within 100 nm distances from sarcomere boundary organelles in Cardiac, Early Postnatal, and Late Postnatal muscles.

Based on the cell type dependence of intrasarcomere CSA heterogeneity demonstrated above (**Figure 1**) and the strong correlation with mitochondrial network orientation (**Figure 2**), we hypothesized that myosin filament curvature would also be greatest in cell types with large proportions of perpendicularly oriented mitochondrial networks. Indeed, while myosin filaments located centrally within a sarcomere were more linear than peripheral myofilaments in mouse cardiac, early postnatal, and late postnatal muscles (**Supplementary Figure 2e**) in addition to the mature slow oxidative and fast oxidative muscles (**Figure 3e**), the magnitude of myosin filament curvature was greater in the fast glycolytic, slow oxidative, and late postnatal muscles compared to the cardiac or early postnatal muscles (**Figure 3g**). Similarly, while proximity to mitochondria and lipid droplets was associated with greater myosin filament curvature than proximity to the SR/T or overall sarcomere boundary for all cell types (**Figure 3h**, **Supplementary Figure 2e-j**), the magnitude of myosin curvature for filaments nearby the larger organelles was also largely dependent on cell type (**Figure 3h**, **Supplementary Figure 2h-i**). Together, these data show that myosin curvature is highest in cell types with perpendicularly oriented mitochondrial networks, while within a given cell, myosin curvature is greater for filaments near mitochondria and lipid droplets compared to the SR/T. Thus, the presence of mitochondria adjacent to a sarcomere, particularly when oriented perpendicular to the contractile axis, influences the shape of the internal myofilaments within that sarcomere.

### Myosin-to-Myosin Lattice Spacing is Variable Along the Sarcomere Length

Force production within a sarcomere is based on physical interactions between actin and myosin, and, thus, is determined in part by the distances between myofilaments^38,39^. Within the A-band of the sarcomere, both actin and myosin are arranged in hexagonal lattice arrays where each myosin filament is surrounded by six other myosin filaments and also by six actin filaments which are closer to each myosin filament and form a smaller lattice^40,41^. Spacing within the myofilament lattices regulates sarcomere shortening velocity^42^, length-tension relationships^33^, cross-bridge kinetics^43,44^, and advective-diffusive metabolite transport^45^ with as small as a 1 nm lattice spacing change correlating with force production^46^. Thus, to better understand the potential functional implications of heterogeneous myofilament structures within a given sarcomere, we determined whether the lattice spacing between myosin filaments was variable among different regions of the sarcomere. Initial observations of single sarcomeres revealed the classic hexagonal myosin lattice within our FIB-SEM datasets although the spacing appeared to vary along the sarcomere length (**Figure 4a-p**, **Supplementary Video 11**). To quantify myosin center-to-center lattice spacing, we performed a 2D fast fourier transform (FFT) analysis of the myosin filament cross-sections at different regions along the length of the sarcomere. Whereas 2D FFT images of single sarcomeres show six bright spots surrounding the image center representing the hexagonal lattice (**Supplementary Video 12**), performing a 2D FFT on the entire cross-section of a muscle dataset results in a circular profile around the center due to the different orientations of the many sarcomeres within the field of view (**Supplementary Figure 3**). To assess lattice spacing in different regions of the sarcomere, we performed whole dataset FFT analyses after segmenting each myosin filament into separate 50 nm regions representing the filament centers, filament ends, and 2-4 intermediate points in between based on their distances from the Z-disk (**Supplementary Figure 3c,f**). In early postnatal muscle, myosin lattice spacing varied by less than 1 nm along the sarcomere length with a 45.91±0.11 nm lattice near the filament ends (n=89 muscle cross-sections) and slightly larger 46.41±0.09 nm spacing near the filament centers (n=88) (**Figure 4q**). Conversely, myosin lattice spacing in late postnatal, fast glycolytic, and slow oxidative muscles all varied by greater than 1 nm along the sarcomere length with larger spacing among the filament centers than for the filament ends (**Figure 4q**). The late postnatal muscle myosin lattice was the most variable of the muscles assessed here and changed by almost 4 nm from 44.28±0.03 nm (n=342) in the sarcomere center to 40.40±0.09 nm (n=333) near the filaments ends. Myosin lattice spacing in the slow oxidative muscles varied by nearly 3 nm from 40.92±0.09 nm (n=138) in the center of the sarcomere to 38.09±0.10 nm (n=93) near the filament ends. Finally, in the fast glycolytic muscles, myosin lattice spacing varied from 41.79±0.07 nm (n=139) near the sarcomere center down to 40.60±0.09 nm (n=137) near the filament ends. These data demonstrate intrasarcomere heterogeneity of myosin-myosin lattice spacing and suggest that the actin-myosin molecular interactions which govern muscle contraction may also vary along the length of a single sarcomere.

**Figure 4:**
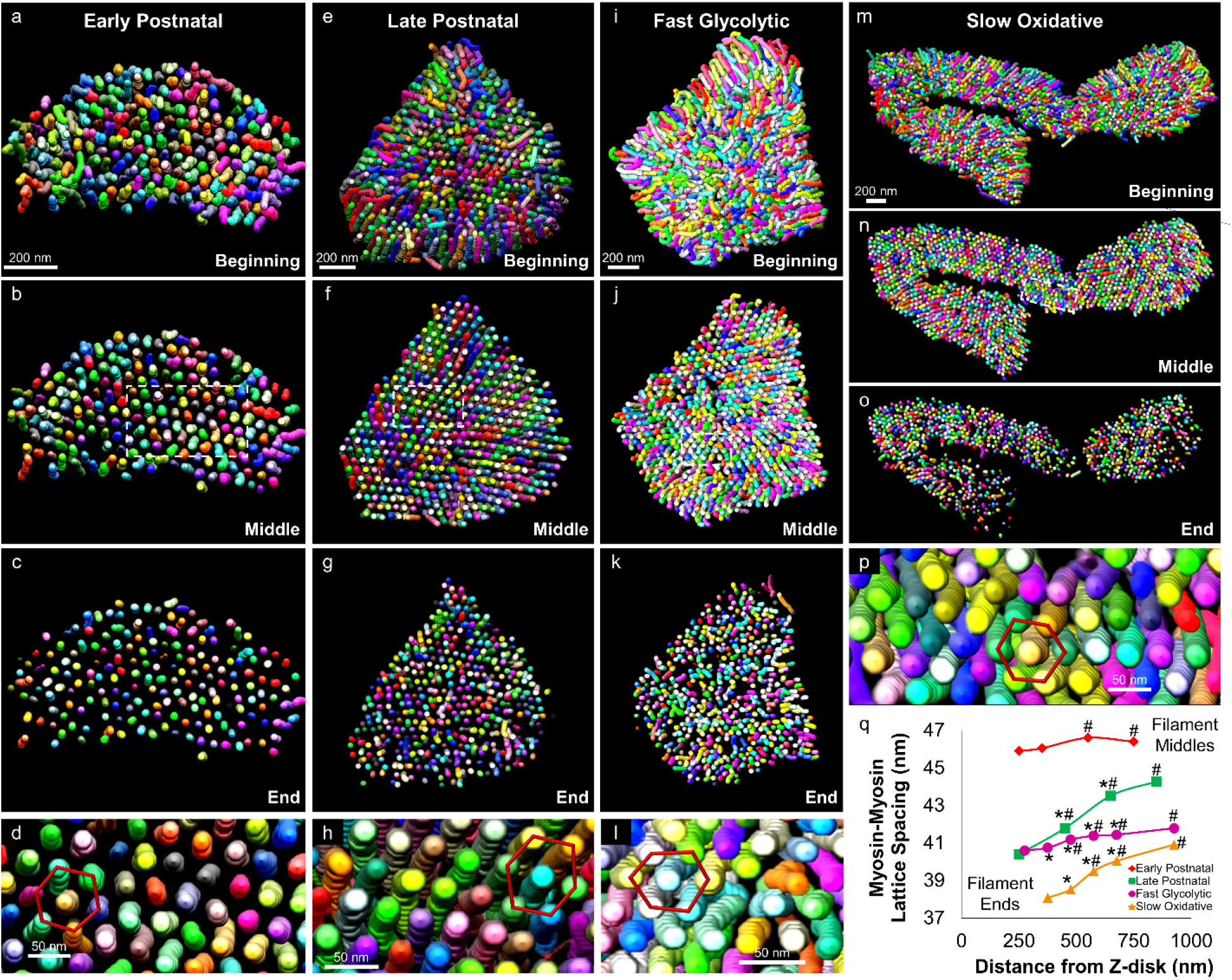
Myosin Lattice Spacing Intrasarcomere Heterogeneity. a-p) 3D renderings of myosin filaments (various colors) within representative single sarcomeres from Early Postnatal (a-d), Late Postnatal (e-h), Fast Glycolytic (i-l), and Slow Oxidative (m-p) muscles showing the entire sarcomere (a,e,i,m), half the sarcomere (b,f,j,n), the ends of the sarcomere (c,g,k,o), and the hexagonal filament lattice from the middle of the sarcomere (d,h,l,p). q) Myosin-myosin lattice spacing as a function of distance from the Z-disk for Early Postnatal (means shown as diamonds), Late Postnatal (squares), Fast Glycolytic (circles), and Slow Oxidative (triangles) muscles. N values: Early Postnatal-88 to 89 muscle cross-sections; Late Postnatal-333 to 342 muscle cross-sections; Fast Glycolytic-137 to 139 muscle cross-sections; Slow Oxidative-93 to 138 muscle cross-sections. Standard error bars are smaller than mean symbols, thus not visible. *significantly different from filament middles. #significantly different from filament ends.

**Supplementary Figure 3:**
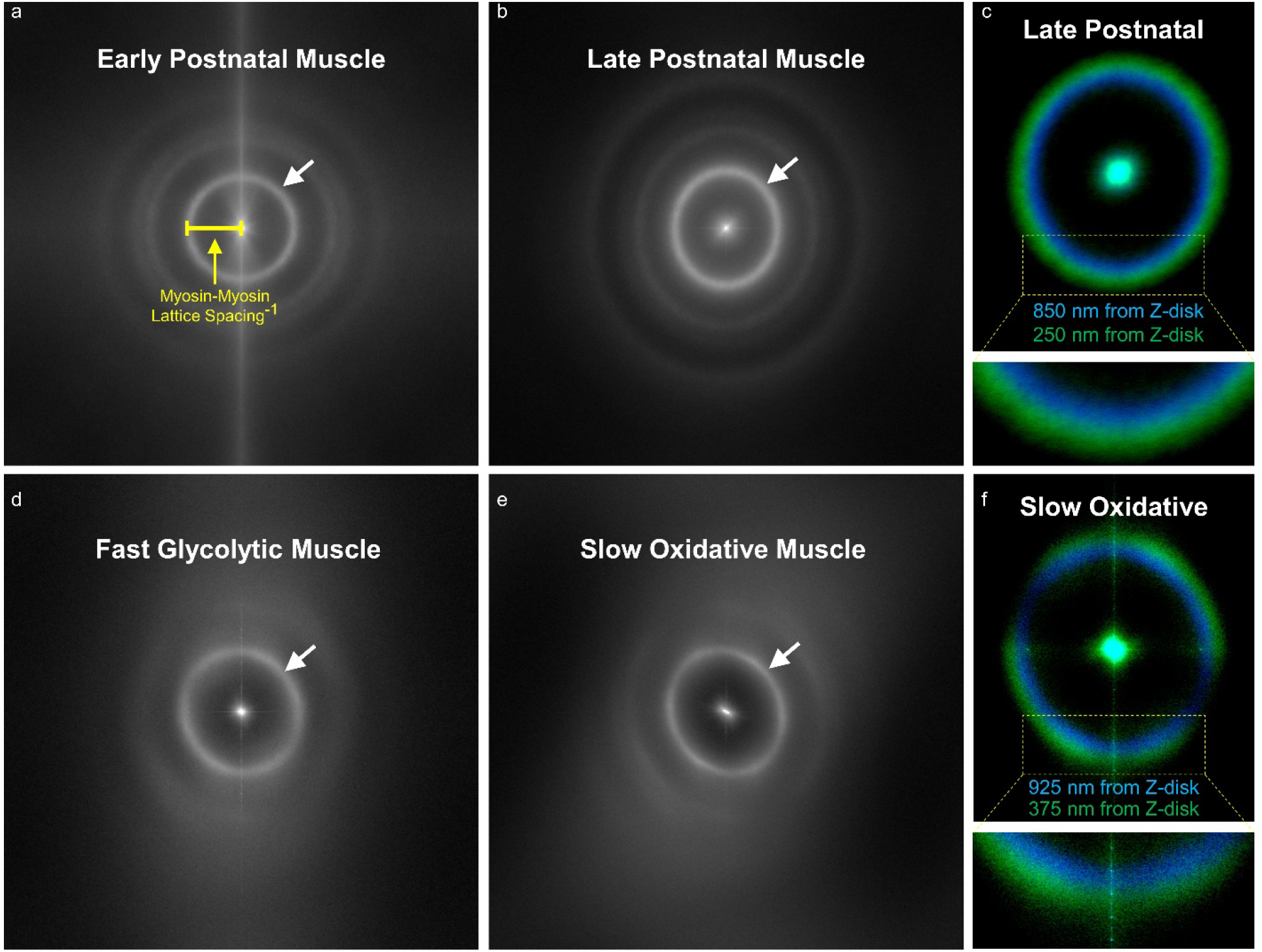
2D FFT Analyses Reveal Intrasarcomere Lattice Spacing Heterogeneity. a,b,d,e) Average of all 2D fast fourier transform (FFT) power spectrums of myosin filament center cross-section images for Early Postnatal (a), Late Postnatal (b), Fast Glycolytic (d) and Slow Oxidative (e) muscles. c,f) Average 2D FFT power spectrum of Late Postnatal myosin filament centers (blue) and ends (green).

## Discussion

The functional benefits of increasing mitochondrial content within a cell are generally well understood to increase the capacity for energy conversion, calcium buffering, ROS/metabolite signaling, and/or other processes in which mitochondria support cellular function^16,47–55^. However, in highly packed cells with limited cytosolic space, such as striated muscle cells, increasing the amount of mitochondria also requires a proportional loss of volume of other cellular structures. As such, there is a functional cost to adding a mitochondrion to a cell that is particularly important to consider when designing therapeutics which may increase mitochondrial content^56–59^. Here, we begin to investigate how striated muscle cells deal with this cost by evaluating how the structure of the sarcomere is affected by proximity to mitochondria across eleven total cell types from three species with a more than 18-fold range in mitochondrial content. We find that the CSA of the sarcomere can be variable along its length with the Z-disk being smaller than the middle of the A-band (**Figure 1**), and that the magnitude of this intrasarcomere heterogeneity across cell types strongly correlates with the proportion of mitochondria located adjacent to the Z-disk (**Figure 2**). These data suggest that placement of a mitochondrion oriented perpendicular to the contractile apparatus comes at the expense of sarcomere CSA, but that the functional cost may be minimized by limiting the reduction in CSA to near the Z-disk rather than the entire sarcomere (**Figure 2o**).

By performing a massively parallel segmentation and analysis of over 2 million myosin filaments (**Figure 3**), we show that intrasarcomere CSA heterogeneity is achieved, at least in part, by the curvature of myosin filaments near the periphery, but not the core, of the sarcomere. Additionally, we find that myosin filaments in close proximity to large intrafibrillar organelles, such as mitochondria, have relatively greater curvature than filaments near smaller diameter organelles such as the sarcoplasmic reticulum. In turn, this variability among myosin filament structures within individual sarcomeres results in reduced lattice spacing near the filament ends compared to the filament centers (**Figure 4**). Alterations in myofilament lattice spacing throughout the entire sarcomere are known to directly affect several measures related to force production (shortening velocity, length-tension relationships, cross-bridge kinetics, etc.)^41–46^. However, it remains unclear how variable lattice spacing within different regions of a single sarcomere influences force production. Moreover, while sarcomeric molecular dynamics simulations of linear actin and myosin filaments have suggested there is an optimal lattice spacing for force production^33^, it is unknown how the different regions within the variable myosin lattices reported here compare to the theoretical optimal spacing. As such, whether smaller lattice spacing closer to the filament ends results in increased force production relative to near the filament centers, perhaps due to increased probability of actin-myosin interactions^60,61^ or potential stress induced activation of myosin in the compressed regions^62,63^, or whether smaller lattice spacing leads to reduced force production, possibly by altering cross-bridge kinetics^43,44,64^, is yet to be determined. With the current absence of methods to directly assess contractile forces with subsarcomeric precision, development of molecular dynamics models accounting for the variability in myofilament spatial relationships along the length of the sarcomere may provide key insights regarding the functional implications of the intrasarcomere structural heterogeneities described here.

Variable lattice spacing within a sarcomere has been suggested to be a consequence of the transformation between the rhombic lattice at the Z-disk and the hexagonal lattice at the M-line^1^. This lattice mismatch would thereby result in oblique angles of the actin filaments and outward radial forces on the Z-disk during contraction which was proposed to be the cause of myofibril splitting during muscle growth^1,22^ However, comparing the intrasarcomere lattice spacing measurements from the four muscle types in **Figure 4** here with our previous assessment of myofibril splitting in the same muscle types^3^ does not support this hypothesis as there is no direct positive or negative relationship between the two measurements (early postnatal muscle: 0.5nm intrasarcomere lattice spacing variability, 27.6% of sarcomeres branch; late postnatal muscle: 3.8nm, 12.8%; fast glycolytic muscle: 1.2nm, 17.5%; slow oxidative muscle: 2.8nm, 43.1%). Thus, while the heterogeneity in intrasarcomere myosin lattice spacing may well be the result of a mismatch between the M-line and Z-disk lattices, intrasarcomere variability in lattice spacing does not appear to be a cause of myofibril splitting.

In summary, by evaluating the 3D relationships among sarcomeres and their adjacent organelles across eleven muscle types from invertebrates to humans, we propose that where a mitochondrion is placed within the intramyofibrillar space influences the structure of the adjacent sarcomeres and can lead to heterogeneity of the cross-sectional area across individual sarcomeres in a cell-type specific manner. In turn, intrasarcomere CSA heterogeneity occurs together with curvature of the internal myosin filaments located at the periphery of the sarcomere with greater curvature of myofilaments located near mitochondria. This intrasarcomere heterogeneity in myosin filament shape thus results in variable lattice spacing among myosin filaments in different regions of the sarcomere. While we are unable to clearly resolve actin filaments in our FIB-SEM datasets, it is likely that the intrasarcomere variability in myosin-myosin lattice spacing occurs concurrently with altered actin-myosin spacing, and thus, the force generating molecular interactions among actin and myosin are also likely variable at different regions within a sarcomere. While it is possible that the chemical fixation procedures used here alter the lattice spacings observed *in vivo*^65–67^, it does not seem likely that fixation would alter the lattice spacing variably in different regions of the sarcomere and in a cell type-dependent manner. Indeed, the *in vivo* immersion fixation procedures used here maintain the circular profiles of muscle capillaries^68^ as well as the mitochondrial diameters observed in live cells^69^ suggesting that the structures reported here are similar to the *in vivo* physiological state. Moreover, curved peripheral myofilaments, including actin, are also observed in cryo-preserved, non-chemically fixed sarcomeres^37^ indicating that the structural heterogeneities within sarcomeres reported here are not simply artifacts of fixation. Thus, overall, these data indicate that the placement of a mitochondrion adjacent to the sarcomere not only alters the energetic support for muscle contraction but also influences the structure of the sarcomere down to the molecular interactions among myofilaments.

## Methods

### Mice

All procedures were approved by the National Heart, Lung, and Blood Institute Animal Care and Use Committee and performed in accordance with the guidelines described in the Animal Care and Welfare Act (7 USC 2142 § 13). 6-8 week old C57BL6/N mice were purchased from Taconic Biosciences (Rensselaer, NY) and fed ad libitum on a 12-hour light, 12-hour dark cycle at 20-26° C. Breeding pairs were setup, and progeny were randomly selected for each experimental group. Early Postnatal mice were from postnatal day 1 (P1) and Late Postnatal mice were from P14. Adult mice were 2-4 months of age. Animals were given free access to food and water and pups were weaned at P21. Due to difficulty using anogenital distance to reliably determine gender in P1 pups, we did not group mice depending on sex, but randomly used both male and female mice.

### Mouse Muscle Preparation

Mouse hindlimb and cardiac muscles were fixed and prepared for imaging as described previously^11^. Mice were placed on a water circulating heated bed and anesthetized via continuous inhalation of 2% isoflurane through a nose cone. Hair and skin were removed from the hindlimbs, and the legs were immersed in fixative containing 2% glutaraldehyde in 100 mM phosphate buffer, pH 7.2 *in vivo* for 30 minutes. For heart fixation, the chest cavity was opened, and cardiac tissue was perfused through the apex of the left ventricle by slowly injecting 2 ml of relaxation buffer (80 mM potassium acetate, 10 mM potassium phosphate, 5 mM EGTA, pH 7.2) followed by 2 ml of fixative solution (2.5% glutaraldehyde, 1% formaldehyde, 120 mM sodium cacodylate, pH 7.2-7.4) through a syringe attached to a 30G needle. After initial fixation, the gastrocnemius, soleus, and/or left ventricles were then removed, cut into 1 mm^3^ cubes, and placed into fixative solution for one hour. After washing with 100 mM cacodylate buffer five times for three minutes at room temperature, the samples were placed in 4% aqueous osmium (3% potassium ferrocyanide, and 200 mM cacodylate) on ice for one hour. The samples were then washed in bi-distilled H_2_O five times for three minutes, and incubated for 20 minutes in fresh thiocarbohydrazide solution at room temperature. Afterwards, the samples were incubated for 30 minutes on ice in 2% osmium solution and then washed in bi-distilled H_2_O five times for three minutes. Next, after incubating in 1% uranyl acetate solution overnight at 4 °C, the samples were washed in bi-distilled H_2_O five times for three minutes and then incubated in lead aspartate solution (20 mM lead nitrate 30 mM aspartic acid, pH 5.5) at 60°C for 20 minutes. After washing in bi-distilled H_2_O at room temperature five times for three minutes, the samples were incubated sequentially in 20%, 50%, 70%, 90%, 95%, 100%, and 100% ethanol for five minutes each, and then incubated in 50% Epon solution and in 75% Epon solution for 3-4 hours and overnight at room temperature, respectively. Epon solution was prepared as a mixture of four components: 11.1 ml Embed812 resin, 6.19 ml DDSA, 6.25 ml NMA, and 0.325 ml DMP-30. Next, the samples were incubated in fresh 100% Epon for one, one, and four hours, sequentially. After removing excess resin using filter paper, the samples were placed on aluminum ZEISS SEM Mounts in 60°C for 2-3 days. Then, using a Leica UCT Ultramicrotome (Leica Microsystems Inc., USA) that is equipped with Trimtool 45 diamond knife, the samples were trimmed, and then gold-coated using a sputter.

### Drosophila Stocks and Muscle Preparation

W flies from the Bloomington Drosophila Stock Center were crossed on yeast corn medium (Bloomington Recipe) at 22° C. Muscles from 2-3 days old flies were dissected on standard fixative solution (2.5% glutaraldehyde, 1% formaldehyde, and 0.1 M sodium cacodylate buffer, pH 7.2) and processed for FIB-SEM imaging as described above.

### Focused Ion Beam Scanning Electron Microscopy

Focus ion beam scanning electron microscopy (FIB-SEM) images were acquired using a ZEISS Crossbeam 540 with ZEISS Atlas 5 software (Carl Zeiss Microscopy GmbH, Jena, Germany) and collected using an in-column energy selective backscatter with filtering grid to reject unwanted secondary electrons and backscatter electrons up to a voltage of 1.5 keV at the working distance of 5.01 mm with a pixel size of 10 nm. FIB milling was performed at 30 keV, 2-2.5 nA beam current, and 10 nm thickness. Image stacks within a volume were aligned using Atlas 5 software (Fibics Incorporated) and exported as TIFF files for analysis. Some raw FIB-SEM datasets were previously used to assess mitochondria-organelle interactions^11,26^, myofibril network connectivity^3,4^, and/or endothelial cell/myocyte interations^68^.

### Image Segmentation

Raw FIB-SEM image volumes were rotated in 3D so that the XY images within the volume were of the muscle cell cross-section. For segmentation of muscle mitochondria, lipid droplets, sarcotubular networks (SRT, sarcoplasmic reticulum + t-tubules), Z-disks, and A-bands, datasets were binned to 20 nm isotropic voxels, and semi-automated machine learning segmentation was performed using the Pixel Classification module in Ilastik^30^ as described previously^11^. Segmentation probability files were exported as 8-bit HDF5 files for import into ImageJ for subsequent analyses. Segmentation of individual mitochondria was performed by using Pixel Classification in Ilastik to generate outer mitochondrial membrane probabilities which were then loaded into the Multicut module in Ilastik for individual organelle separation as described previously^11^.

For segmentation of individual myosin filaments, raw FIB-SEM data were upscaled to 5 nm isotropic voxels using bicubic interpolation within the Scale feature in ImageJ^70^ resulting in 8-bit datasets of up to 160 Gb. A series of raw data images were loaded into complete volume memory in Thermo Scientific Avizo Software 2020.3 with the XFiber extension (Thermo Fisher Scientific, Waltham, MA) making sure to specify voxels were 5 nm. The XFiber extension permits segmentation of tightly packed cylindrical objects by first computing normalized cross correlation of the images against a hollow cylindrical template and by then tracing centerlines along filaments^71^.

Cylinder correlation was performed using a template cylinder length of 100 nm, angular sampling of 5, mask cylinder radius of 20, outer cylinder radius of 19, and an inner cylinder radius of 0. Using a Windows 10 desktop PC with 64 logical processors (Intel Xeon Gold 6142M), 2.0 TB of RAM, and an NVIDIA Quadro RTX8000 48 Gb GPU, cylinder correlation took 3-12 days depending on the size of the dataset and parameters chosen. Myosin filaments were then segmented from the resulting correlation and orientation fields using the Trace Correlation Lines module in Avizo. Minimum Seed Correlation (range 105-125) and Minimum Continuation (range 60-95) values within the Trace Correlation Lines module were varied per dataset using inspection of the correlation field images as guide for correlation values which best corresponded to myosin filaments in the raw data. Direction coefficient was set to 0.3, Minimum Distance was 20, Minimum Length was 500 nm, Search Cone Length, Angle, and Minimum Step Size were 100 nm, 30, and 10%, respectively. Trace Correlation Lines typically took 0.5 – 3 days depending on dataset size and parameters chosen. The resultant correlation lines were then converted with the Convert Geometry to Label module using the input raw dataset size parameters and saved as a 3D .raw file for import into ImageJ for subsequent analyses.

### Image Analysis

Ilastik 8-bit HDF5 probability files for organelle and contractile structures were imported into ImageJ using the Ilastik plugin and made into binary files by using an intensity threshold of 128. Small segmentation errors were filtered out of the binary image volumes using the Remove Outliers plugin with block radius values from 3-10 and standard deviation values from 1.5 to 2.0. Z-disk CSA was assessed per half sarcomere sheet (**Supplementary Movie 4**) by first performing a Grouped Z maximum projection (Image-Stacks-Tools) at the predetermined sarcomere length. Because each Z-disk is part of two adjacent half sarcomeres, the maximum projection image was duplicated and the two images interleaved (Image-Stacks-Tools) together. Then the intensity of each image assessed using Plot Z-axis profile (Image-Stacks) and the data copied and pasted into Excel (2016, Microsoft) for comparison to the corresponding A-band CSA values. A-band half sarcomere sheet CSA was assessed by first performing a Grouped Z average projection at half sarcomere length. Binary images were created from the average projection by normalizing local contrast (Plugins-Integral Image Filters) with a quarter sarcomere radius, 10 standard deviations, and marking the stretch and center boxes. The resultant image was masked by the A-band Grouped Z maximum projection at half sarcomere length and then intensity thresholded using the auto default value.

Plot Z-axis profile was then used to assess the A-band intensity for each image, and the data copied and pasted to Excel where it was divided by the corresponding Z-disk intensity value yielding a relative CSA difference per half sarcomere sheet.

Z-disk adjacent mitochondrial abundance was determined by first separating the mitochondrial networks into two regions based on the proximity to the Z-disk. A Distance Transform 3D (Plugins-Process) was performed on the binary Z-disk image volumes and thresholded at a value 200 nm larger than the width of the I-band visible in the raw FIB-SEM datasets. This value was determined to encompass all perpendicularly oriented mitochondria based on analyses of the late postnatal and mature muscles in the mouse, leg muscles in *Drosophila,* and human muscles. A mask of the Z-adjacent regions was then created by subtracting (Process-Math) 254 from the thresholded Z-disk distance transform image, and the mask was multiplied by the binary mitochondrial segmentation image to create a Z-adjacent mitochondrial image volume. The total number of Z-adjacent mitochondrial pixels was determined from a Histogram (Analyze) of the entire stack and divided by the total number of mitochondrial pixels from the original binary mitochondrial image. Total mitochondrial content was determined by dividing the total number of mitochondrial pixels by the total number of cellular pixels.

For myosin filament analyses, the .raw file from Avizo was imported into ImageJ (File-Import) making sure to enter the corresponding Width and Height pixel values and Number of images. Myosin filament deviation from linearity was assessed using the Particle Analyser within the BoneJ^72,73^ plugin for ImageJ (Plugins-BoneJ-Analyze) and selecting only the Moments of Inertia and Show Particle Stack options. The resultant data table and labeled filament image volume were saved for further analyses. The moment of inertia along the longest principal axis (I3) was then divided by the volume for each myosin filament to determine its deviation from linearity^74^ by using a custom ImageJ macro to perform math operations within results tables. Visualization of filament deviation from linearity values was performed by using the Assign Measure to Label function within the MorphoLibJ^75^ plugin (Plugins-MorphoLibJ-Label Images) using the labeled filament image and the I3/volume values from the results table.

For comparisons of filament deviation from linearity based on organelle proximity, image volumes of binary organelles segmented with 20 nm voxels were first upscaled (Image-Scale) to 5 nm voxels without interpolation. A Distance Transform 3D (Plugins-Process) was then performed on the mitochondria, lipid droplet, SRT, and total sarcomere boundary (mitochondria+lipid droplet+SRT) binary image volumes. The minimum distance between each myosin filament and the muscle organelles was then determined using the Intensity Measurements 2D/3D module within the MorphoLibJ plugin selecting the respective organelle distance transform image as Input, the myosin filament label image as Labels, and selecting only the Min Measurements box. The resultant tables were then saved and appended to the corresponding labels in the I3/volume table from above in Excel (for up to 1,048,576 labels) or SPSS (for more than 1,048,576 labels). Filaments that were greater than 1 μm in length and did not overlap with any organelle were used for all proximity analyses.

Myosin filament center-to-center distances were assessed by 2D FFT analyses (Process-FFT) at different regions along the filament length. The myosin filament .raw file was imported into ImageJ as above and then multiplied (Process-Image Calculator) by the Z-disk distance transform image. The resultant filament distance from Z-disk image was then thresholded (Image-Adjust-Manual Threshold) to select a 50 nm region of the filament centers, near the filament ends, or at intermediate points in between and made into a binary image (Process-Binary-Make Binary). A maximum Z-projection (Image-Stacks-Tools-Grouped Z Project) was then performed for every 50 nm of the resultant binary image, and a 2D FFT was run for each projection image using a custom ImageJ macro to assess the periodicity of the myosin to myosin distances. The intensity of each resultant 2D FFT image was assessed using a custom ImageJ macro to iteratively select all pixels of a given distance from the FFT center using a distance map from the FFT center (Plugins-Process-Exact Euclidean Distance Transform) and measuring their intensity (Plugins-MorphoLibJ-Analyze-Intensity Measurements 2D/3D). The resultant intensity profiles for each 2D FFT were then loaded into Excel, and the maximum intensity value for all distances between 30 and 60 nm was selected as the myosin to myosin filament distance for each respective image.

### Geometric Model of Sarcomere Isometric Force Production

Of the two simulated scenarios, the first scenario is a uniform CSA reduction with a proportional reduction of the number of myofilaments, keeping the lattice spacing unchanged. Assuming that the isometric force production of each filament also remains unchanged, then the total force is proportional to the total number of filaments. A reduction in the CSA directly results in an identical reduction in the isometric force. The second scenario simulates a myofibril of a circular cross-section, where the CSA tapers down gradually from the sarcomere center to the Z-disk in a smooth arc shape. It was assumed that the lattice spacing tapers down uniformly, and the tensile force of individual myofilaments was not altered by the compression. Each filament “pulled” on the Z-disk at a slightly off-perpendicular angle due to the variable curvature of the filament. This lead to a reduction of the axial force according to the sine factor of the filament insertion angle at the Z-disk. The overall effect on the force production was analytically derived by the integral of the axial force from each filament within the sarcomere bundle.

### Image Rendering

Movies of 3D renderings of organelle and contractile structures were generated in Imaris 9.5 (Bitplane). Pictures of 3D renderings were created either using Imaris or the Volume Viewer plugin in ImageJ.

### Statistical Analysis

Quantitative data was assessed using Excel 2016 (Microsoft), Prism 9.0.0 (Graphpad), or SPSS 28.0.0.0 (IBM) for all statistical analyses. All comparisons of means were performed using a one-way ANOVA with a Tukey’s HSD post hoc test. Linear regression analyses were performed in SPSS (Analyze-Regression-Linear) using default settings. A p-value < 0.05 was used to determine statistical significance.

## Data Availability

The raw FIB-SEM datasets generated and/or analysed during the current study are available at (https://doi.org/10.5281/zenodo.5796264).

## Acknowledgements

We would like to acknowledge Eric Lindberg and Ye Sun for assistance with collecting FIB-SEM datasets. We thank Brian Caffrey (University of British Columbia), Sriram Subramaniam (University of British Columbia), and Luigi Ferrucci (National Institute of Aging) for providing access to the raw human muscle FIB-SEM datasets from the Baltimore Longitudinal Study of Aging. We are also grateful to Kenneth Campbell, Thomas Irving, James Sellers, Neil Billington, Kenneth Taylor, and Malcolm Irving for helpful discussions regarding the functional implications of the structures described here. This work was supported by the Division of Intramural Research of the National Heart Lung and Blood Institute and the Intramural Research Program of the National Institute of Arthritis and Musculoskeletal and Skin Diseases.

## Author Contributions

PK, BG and YK prepped tissues for imaging. PK, BG, YK, and CKEB designed and CKEB performed imaging experiments. ASH, PTA, TBW, and BG designed and performed image analysis and BG created figures and videos. HW designed and performed computational modeling experiments. BG wrote and PK, ASH, PTA, YK, TBW, HW, CKEB, and BG edited and approved the manuscript.

## Materials and Correspondence

Further information and requests for resources and reagents should be directed to Brian Glancy.

## Supplementary Video Legends

**Supplementary Video 1:** Cross-sectional view of three serial sarcomeres from two parallel myofibrillar segments from a fast twitch glycolytic adult mouse gastrocnemius muscle visualized by focused ion beam scanning electron microscopy with 6 nm isotropic pixel size.

**Supplementary Video 2:** 3D rendering and rotation of the adult gastrocnemius muscle sarcomeres (translucent yellow and cyan), Z-disks (red), mitochondria (magenta), sarcoplasmic reticulum (green), and t-tubules (blue) from the raw data shown in Supplementary Video 1.

**Supplementary Video 3:** 3D rendering and rotation of sarcomeres (translucent yellow and cyan), Z-disks (red), and mitochondria (magenta) from *Drosophila* indirect flight muscle.

**Supplementary Video 4:** 3D rendering and rotation of the Z-disk (light green and various colors) and sarcomere (magenta) sheets that comprise the *Drosophila* jump muscle.

**Supplementary Video 5:** 3D rendering and fly through of the mitochondrial networks in *Drosophila* indirect flight (green), direct flight (blue), jump (yellow), and leg (magenta) muscles.

**Supplementary Video 6:** 3D rendering and fly through of the grid-like mitochondrial network of a slow-twitch oxidative mouse gastrocnemius muscle. Raw FIB-SEM data (greyscale) is pulled back to reveal the mitochondrial network segmented into Z-disk adjacent (light blue) and non-Z-disk adjacent (red) mitochondria.

**Supplementary Video 7:** 3D rendering and fly through of the parallel mitochondrial network of a mouse cardiac muscle. The mitochondrial network is segmented into Z-disk adjacent (light blue) and non-Z-disk adjacent (red) mitochondria.

**Supplementary Video 8:** 3D rendering and fly through of all the myosin filaments (various colors) from a mouse late postnatal soleus muscle FIB-SEM dataset.

**Supplementary Video 9:** 3D rendering and fly through of all the myosin filaments from a mouse late postnatal soleus muscle FIB-SEM dataset. Filaments are colored according to their deviation from linearity where more linear filaments are dark blue and more curved filaments are yellow/white.

**Supplementary Video 10:** 3D rendering and rotation of the myosin filaments within a single mouse cardiac sarcomere. Adjacent mitochondria (red), sarcotubular network (green), and a lipid droplet (cyan) are also shown.

**Supplementary Video 11:** 3D rendering and fly through of the myosin filaments (various colors) within a single mouse slow-twitch oxidative muscle sarcomere.

**Supplementary Video 12:** Cross-sectional view of the segmented myosin filaments (white) along the length of a single sarcomere and the corresponding 2D fast fourier transform power spectrum from early postnatal, late postnatal, slow-twitch oxidative, and fast-twitch glycolytic muscles.

